# A comprehensive workflow for allele-specific immune gene quantification and expression analysis in single-cell RNA-seq data

**DOI:** 10.1101/2024.12.10.627679

**Authors:** Ahmad Al Ajami, Jonas Schuck, Federico Marini, Katharina Imkeller

## Abstract

**Motivation:** Immune molecules such as B and T cell receptors, human leukocyte antigens (HLAs), or killer Ig-like receptors (KIRs) are encoded in the most genetically diverse loci of the human genome. Many of these immune genes exhibit remarkable allelic diversity across populations. While computational methods for HLA typing from bulk RNA sequencing data have emerged, streamlined solutions for allele-specific quantification in single-cell RNA sequencing (scRNA-seq) are lacking. Moreover, no standardized data structure or analytical framework has been established to handle allele-specific immune gene expression data at single-cell level.

**Results:** We present a comprehensive workflow to (1) automate allele-typing and allele-specific expression quantification of HLA transcripts in scRNA-seq data using a Snakemake workflow, *scIGD* (single-cell ImmunoGenomic Diversity), and (2) represent and interactively explore immune gene expression at different annotation levels using a multi-layer data structure implemented as an R/Bioconductor software package, *SingleCellAlleleExperiment*. We validated our approach on a diverse spectrum of scRNA-seq datasets, and found that it performs consistently across different sequencing platforms and experimental setups. We illustrate how our method can be utilized to study loss of HLA expression in tumor cells or discover differential HLA allele expression in specific immune cell subtypes. By capturing such allele-specific expression patterns and their variation, our workflow offers novel insights into human immunogenomic diversity.

**Availability and implementation:** *scIGD* is available under the MIT license at: https://github.com/AGImkeller/scIGD. *SingleCellAlleleExperiment* is available under the MIT license at: https://bioconductor.org/packages/SingleCellAlleleExperiment.

*scaeData* provides validation datasets and is available under the MIT license at: https://bioconductor.org/packages/scaeData.

Data processed with *scIGD* are available at: https://doi.org/10.5281/zenodo.14033960.

**Contact:** Katharina Imkeller.

**Supplementary information:** Supplementary data are available within the same submission.

## 1 INTRODUCTION

Immune genes play a critical role in mediating antigen recognition and exhibit a high degree of structural diversity, which is essential for adapting to the vast array of pathogens encountered by the immune system. This diversity is achieved through two primary genetic mechanisms: polygeny and hyperpolymorphism [1]. Polygeny refers to the presence of multiple functionally-similar genes that encode similar protein subunits, as for example *IGLC2* and *IGLC7* genes which both encode an immunoglobulin light chain constant domain. On the other hand, hyperpolymorphism denotes the presence of numerous allelic variants of the same gene across individuals. As a key example of this immunogenomic diversity, we focus on the major histocompatibility complex (MHC) in this study. Also known as human leukocyte antigens (HLAs) in humans, the MHC molecules present antigens to T cells, thereby modulating adaptive immune responses. The HLA region is located on chromosome 6 and includes more than 200 genes [2,3]. This includes three principal class I genes—*HLA-A, HLA-B*, and *HLA-C*—and three pairs of class II genes—*HLA-DR, HLA-DP*, and *HLA-DQ*. These genes exhibit remarkable diversity, with around 28000 HLA class I alleles and approximately 12400 HLA class II alleles identified to date [2,3].

Most individuals are heterozygous for HLA genes, inheriting different alleles from each parent. Codominant expression of HLA alleles, where both alleles are equally expressed, enables individuals to present a broader repertoire of antigens [1]. HLA class I molecules, expressed on nearly all nucleated cells, present antigens to CD8+ T cells, while HLA class II molecules, predominantly found on specialized immune cells like dendritic cells, macrophages, and B cells, present antigens to CD4+ T cells.

With the growing relevance of immunotherapy in cancer treatment, precise HLA typing and quantification of expression levels have become critical because they enable a better understanding of how HLAs present tumor neoantigens on the cell surface [4]. Several tools have therefore been developed for HLA typing from sequencing data, including *seq2HLA* [5], *OptiType* [6], *PHLAT* [7], *HLAProfiler* [8], and *arcasHLA* [9], with benchmarking studies indicating *arcasHLA*’s consistent performance [10,11]. Additionally, tools like *scHLAcount* construct personalized references for HLA allele-specific quantification [12]. However, there remains a significant gap in tools designed to streamline HLA allele typing and quantification specifically for single-cell RNA-seq (scRNA-seq) data. Existing methods lack an integrated, user-friendly pipeline that supports both HLA allele-typing and allele-specific expression quantification, and enables interactive exploration of immune gene expression.

In this work, we develop a comprehensive workflow to bridge the gap in analyzing HLA expression from scRNA-seq data (Fig. 1). Our approach not only automates HLA allele-typing and enables allele-specific quantification from scRNA-seq data using a Snakemake-based [13] workflow, *scIGD* (single-cell ImmunoGenomic Diversity), but also offers a versatile multi-layer data structure implemented in the R/Bioconductor package, *SingleCellAlleleExperiment* (*SCAE*), which allows for the representation of HLA at multiple levels, ranging from alleles to genes and functional classes. *SCAE* builds directly on the *SingleCellExperiment* (*SCE*) [14] class and thereby ensures compatibility with existing Bioconductor tools for single-cell analysis. Example datasets to assist users in testing the workflow are available in our corresponding R/ExperimentHub [15] data package, *scaeData*.

**Fig. 1.**
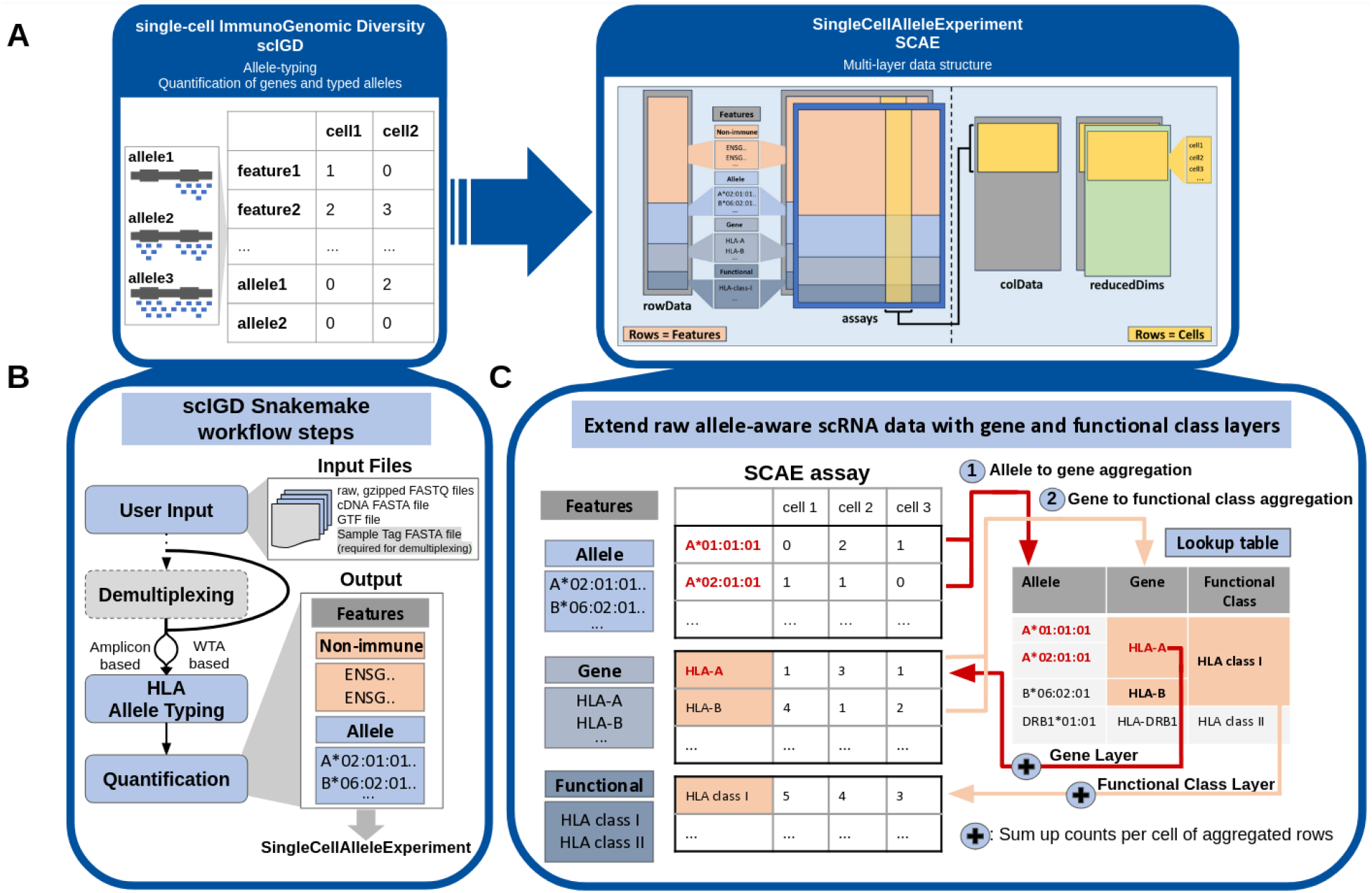
Overview of the workflow. **A)** Overview of the integration between the two key tools in our workflow: the *scIGD* Snakemake workflow and the *SingleCellAlleleExperiment* R/Bioconductor package. **B)** Detailed schematic representation of *scIGD*’s processing steps, illustrating how raw sequencing data is processed to generate HLA allele-typing and quantification results. Non-immune genes are quantified at the conventional gene level, whereas immune gene quantification is performed at the allele level. The output serves as input for downstream analysis in *SingleCellAlleleExperiment*. **C)** Detailed schematic representation of the multi-layered data structure within *SingleCellAlleleExperiment*, showing how HLA data is organized across allele, gene, and functional class layers. With the help of a lookup table, allele counts outputted by *scIGD* and belonging to the same gene are aggregated to populate the gene layer (1). Resulting gene counts belonging to the same functional class (e.g. HLA Class I and Class II) are further aggregated to populate the functional class layer (2).

By capturing HLA allele-specific variations and providing a detailed view of HLA expression, our workflow is able to detect HLA allele-specific expression differences across immune cell subsets, as well as HLA loss in tumor samples. Notably, it enables a more accurate and deeper exploration of immune functions and responses, facilitating data-driven immunological research.

## 2 MATERIALS AND METHODS

### 2.1 Software description

The Snakemake workflow of *scIGD* is compatible with Unix-based systems, while *SingleCellAlleleExperiment* and *scaeData* are platform-independent, supporting Windows, macOS, and Linux. Each tool is optimized for minimal dependencies, ensuring easy installation and use. Comprehensive documentation and usage examples are available under the respective links: scIGD: https://github.com/AGImkeller/scIGD;

SingleCellAlleleExperiment: https://bioconductor.org/packages/SingleCellAlleleExperiment; scaeData: https://bioconductor.org/packages/scaeData.

### 2.2 scIGD

*scIGD* (single-cell ImmunoGenomic Diversity) is a Snakemake-based workflow designed to automate the typing of HLA alleles and enable allele-specific quantification from scRNA-seq data (Fig. 1B). The primary input for *scIGD* consists of raw, gzipped FASTQ files, which are processed through the workflow’s various stages to generate allele-typing and quantification results.

The workflow is organized into three distinct stages (Demultiplexing, HLA allele-typing, and Quantification), each addressing specific objectives.

#### 2.2.1 Demultiplexing

The first stage of the *scIGD* workflow focuses on demultiplexing scRNA-seq datasets which contain reads from multiple donors that were barcoded using sample tags (for example using the BD Single-Cell Multiplexing Kit). This step is omitted if the input data does not contain sample tag information or consists of only a single donor. The primary goal of this step is to generate donor-specific FASTQ files, which are essential for subsequent allele typing.

To perform demultiplexing, the user must provide two key inputs: the raw, gzipped scRNA-seq FASTQ files (R1 and R2) and a sample tag sequences FASTA file, which contains the unique sequence tags corresponding to each sample. Using *kallisto-bustools* [16], the input data is processed to produce a count matrix that links cell barcodes with sample tags. For each cell barcode, the method selects the sample tag ID that accounts for at least 75% of the read count. If no single sample tag meets this threshold, the cell is classified as a multiplet and excluded from further analysis.

After the correct sample tag is assigned to each cell barcode, the workflow matches read IDs with cell barcodes and sample tags and splits the original FASTQ files into multiple donor-specific FASTQ files, one for each sample tag. These donor-specific files are then used as inputs for the next stages of the workflow. Demultiplexing in *scIGD* is currently implemented for scRNA-seq datasets barcoded using the BD Single-Cell Multiplexing Kit.

#### 2.2.2 HLA allele-typing

The second stage of the *scIGD* workflow focuses on HLA allele-typing. Our workflow supports both amplicon-based and whole-transcriptome-based scRNA-seq data.

For amplicon-based scRNA-seq, *scIGD* implements a minimal allele-typing which can only detect allelic differences located in the part of the amplicon that is covered by the respective sequencing approach. A FASTA file containing the cDNA sequences of target amplicons is required as input. For a defined set of amplicons of interest, the workflow infers the one or two most abundant transcript variants that can be distinguished given the implemented sequencing coverage. Since these transcript variants typically do not allow identification of reference alleles, the transcript variants are assigned arbitrary allele identifiers (e.g. *HLA-A-AV-1, HLA-A-AV-2, HLA-A-AV-3*, where AV refers to Allelic Variant).

For whole-transcriptome-based (WTA) scRNA-seq data, a more comprehensive HLA allele-typing is performed using the existing *arcasHLA* methodology [9]. This stage utilizes a reference genome FASTA file and matching GTF file with genomic annotations. Both files should be downloaded from the Ensembl database [17]. The *STAR* aligner [18] is used to map measured sequencing reads to the reference genome. *arcasHLA* then extracts those reads that map to chromosome 6, which contains the HLA loci, and uses the IMGT/HLA database [2,3] and an expectation-maximization (EM) model to determine the combination of HLA alleles that best explains the measured sequencing data.

The allele-typing stage of the *scIGD* workflow generates a donor-specific reference cDNA FASTA file in which the immune gene references are replaced by the cDNA sequences of the previously typed alleles.

#### 2.2.3 Quantification

The final stage of the *scIGD* workflow focuses on the quantification of genes and typed alleles. This stage applies equally to amplicon-based and whole-transcriptome-based scRNA-seq data.

*kallisto-bustools* [16] is used to perform alignment-free quantification of the transcripts represented in the donor-specific cDNA FASTA reference. Importantly, since the donor-specific cDNA reference may contain two alleles of the same gene, and thus very similar sequences, *scIGD* uses the *-mm* option of *kallisto-bustools* that includes reads that pseudo-align to multiple transcripts in the reference.

Finally, a lookup table is created to map each allele to its corresponding gene and functional class. This table is crucial for constructing the *SingleCellAlleleExperiment* data structure used in all subsequent steps, because it defines the relationships between alleles, genes, and their functional classes.

#### 2.2.4 Setting up the scIGD analysis workflow

To use *scIGD*, users must first set up a working directory and create an environment with the required software. We recommend using *conda* or *mamba* for managing the environment, and a detailed step-by-step guide for this process is available in the GitHub repository (https://github.com/AGImkeller/scIGD).

As a Snakemake workflow, *scIGD* includes a configuration file that allows users to adjust the workflow according to their specific experimental setup and data. This config file supports the customization of various parameters of the workflow. Detailed examples and descriptions of these parameters are also provided in the repository to help users with the setup.

The entire *scIGD* workflow can be executed with a single command, offering a streamlined and user-friendly experience. Upon installing *scIGD*, several files integral to the workflow are automatically set up (Fig. 2A), while additional input files must be supplied by the user. An example of the required input data for a WTA scRNA-seq dataset is shown in Fig. 2B. All output files generated by the workflow are stored in a user-defined output directory, along with detailed log files for each step. Fig. 2C provides an overview of the output files produced when *scIGD* is run on a WTA dataset.

**Fig. 2.**
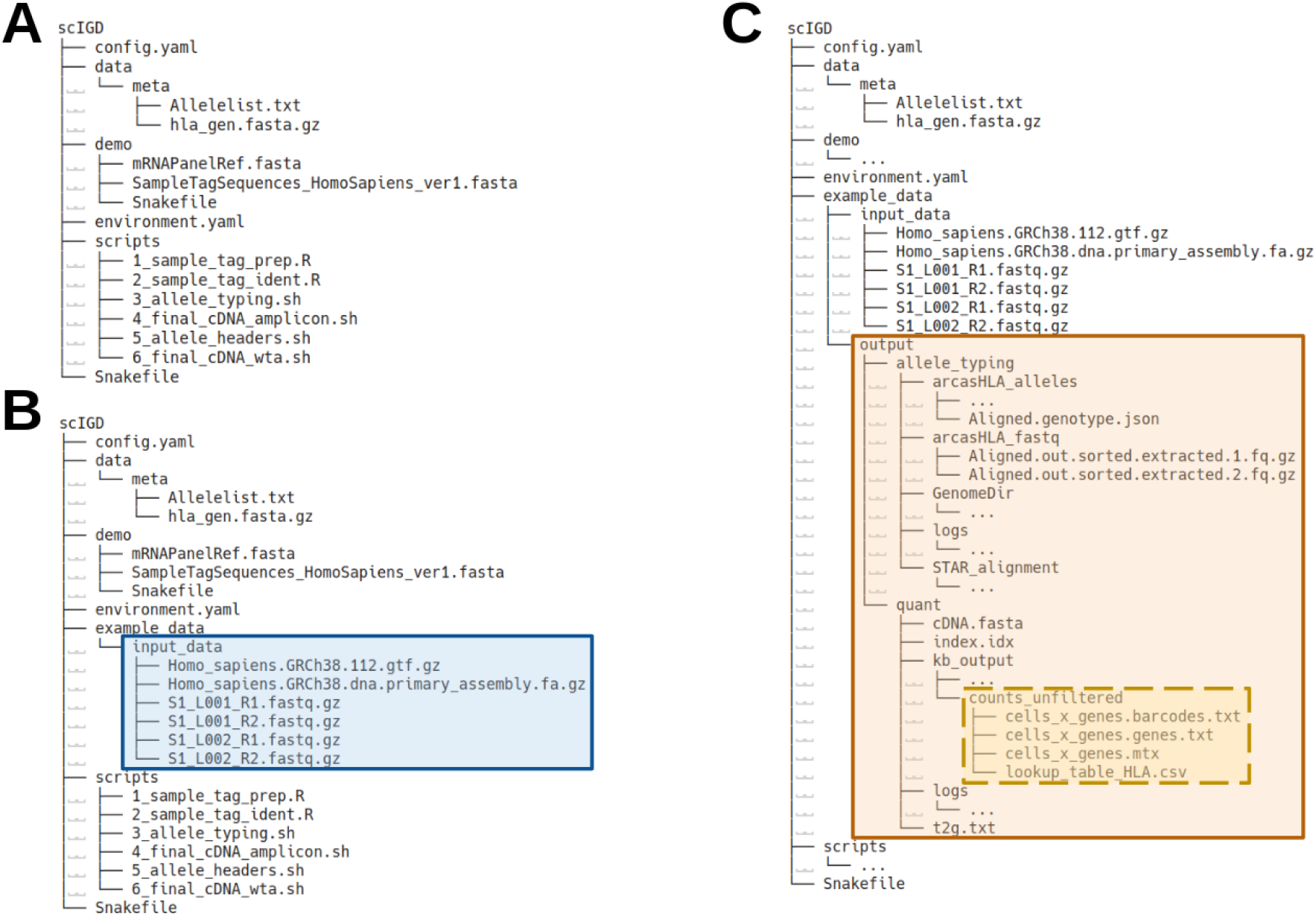
File and directory structure in the scIGD workflow. **A)** File and directory structure provided upon installation of the *scIGD* workflow. **B)** Exemplary directory structure for setting up *scIGD* to analyze a WTA-based dataset. The input files required are highlighted in blue. While this layout serves as a guideline, the specific file paths can be customized within the config.yaml file. **C)** Overview of the output files generated from running *scIGD* on a WTA-based dataset. The output files generated are highlighted in orange. This includes log files for each processing step, as well as the outputs generated by the tools involved in both HLA allele-typing and quantification stages. Directories with “…” indicate that additional files are present but have been collapsed for simplicity. The four key files required for downstream analysis with the *SingleCellAlleleExperiment* package are highlighted in yellow and enclosed within a dashed box.

### 2.3 SingleCellAlleleExperiment

The second component in our workflow is *SingleCellAlleleExperiment* (*SCAE*), an R/Bioconductor package that extends *SingleCellExperiment* (*SCE*) [14] to support immune gene quantification data generated by *scIGD*. By building directly on *SCE, SCAE* inherits its robust data structure and compatibility with a vast array of existing Bioconductor packages for single-cell analysis. This ensures users can seamlessly integrate *SCAE* objects into established workflows without the need to rewrite or duplicate functionality, leveraging the extensive ecosystem of tools already designed for single-cell data.

*SCAE* offers a multi-layer framework within a single data object, allowing users to analyze immune gene expression across different annotation levels. *SCAE* imports the count matrix and lookup table generated by *scIGD*. Following the relationships defined in the lookup table, it aggregates allele-level counts belonging to the same gene into gene-level counts. Resulting gene counts belonging to the same functional class (e.g. HLA Class I and Class II) are further aggregated into functional-level counts (Fig. 1C). Such a multi-layer design provides flexibility, allowing users to focus on specific data layers and adapt analyses as needed, and at the same time makes this object compatible with the wealth of existing packages for single-cell data analysis in the Bioconductor ecosystem. *SCAE* also supports quality control and preprocessing steps like cell filtering by knee plot, and normalization that accounts for the extended feature set. The *scaeData* package provides curated example datasets to serve as a practical resource for users to explore the features and capabilities of the *SingleCellAlleleExperiment* package.

### 2.4 Description of the presented datasets

With the exception of the demultiplexing step, the *scIGD* workflow is compatible with both 10x (different chemistry types are supported) and BD Rhapsody platforms, supporting WTA and amplicon-based scRNA-seq data.

We validated the performance on four different datasets: 14 donors from the AIDA cohort (8 Korean, 6 Japanese, chosen from the complete cohort based on highest sequencing depth) [19], the Merkel-cell carcinoma dataset (PBMC and tumor samples, pre- and post-treatment) [20], the 20k PBMC dataset [21], and a multiplexed Multiple Myeloma dataset with 8 donors [22] (Table 1).

**Table 1.** Summary of scRNA-seq datasets analyzed using the *scIGD* workflow.

| Dataset | Library | Experimental Setup | Multiplexed | Number of Donors | Longitudinal Study | Reference |
| --- | --- | --- | --- | --- | --- | --- |
| AIDA | 10x 5' v2 | WTA | NO | 14 | NO | Kock et al., 2024 |
| Merkel-Cell Carcinoma | 10x 3' v2 | WTA | NO | 1 | YES | Paulson et al., 2018 |
| 20k PBMC | 10x 3' v3 | WTA | NO | 1 | NO | 10x Genomics |
| Multiple Myeloma | BD Rhapsody | Amplicon | YES | 8 | NO | Enssle et al., 2024 |

### 2.5 Preprocessing of datasets after the scIGD quantification

We applied the functionality of *SCAE* and performed downstream analysis on the Merkel-cell carcinoma and 20k PBMC datasets. When creating the *SCAE* objects, we filtered out data from empty droplets using the *filter_mode=“yes”* option, which filters according to the inflection point of a knee plot based on barcode ranks. Additionally, we enabled *log=TRUE*, which normalizes the raw counts, ensuring that multiple layers of HLA features do not disproportionately impact the size factors during normalization. We then performed comprehensive preprocessing, including the exploration of quality control (QC) metrics and subsequent filtering of cells and features for robust downstream analysis.

For the Merkel-cell carcinoma dataset, all four samples were merged into a single *SCAE* object. After batch correction using the mutual nearest neighbors (*MNN*) method [23], we performed principal component analysis (*PCA*) on highly variable genes to reduce the dimensionality of the data while identifying and retaining main sources of variation among the cells. The results were projected onto a t-distributed stochastic neighbor embedding (*t-SNE*) plot. To ensure accurate comparisons across samples, we normalized and logarithmically transformed the counts using *multiBatchNorm*, accounting for the sample information [23].

For the 20k PBMC dataset, raw counts were normalized and logarithmically transformed using *logNormCounts* [24]. *PCA* was performed on the highly variable genes prior to constructing the shared nearest neighbor (*SNN*) graph and performing Louvain clustering with *igraph* [25]. The Louvain clustering result with *k=50* returned a reasonable number of clusters and was used in downstream analysis. Importantly, since *SCAE* builds directly on the *SCE* class, we were able to use existing *SCE*-based functions and methods for all analysis steps mentioned above.

## 3 RESULTS

Our newly developed workflow comprises two software tools: *scIGD*, implemented using Snakemake, and *SingleCellAlleleExperiment*, an R/Bioconductor package (Fig. 1A). *scIGD* is designed as a Snakemake workflow to perform HLA allele-typing and allele-specific quantification from scRNA-seq data (Fig. 1B). On the other hand, *SingleCellAlleleExperiment* is designed to run entirely within an R environment, allowing for multi-layer representations of immune gene quantification data, ranging from allele-to gene- and functional-level annotations (Fig. 1C).

### 3.1 scIGD accurately types and quantifies HLA alleles in a broad range of single-cell sequencing datasets

*scIGD* performs HLA-allele typing to determine the HLA alleles expressed by an individual using *arcasHLA* [9], a tool chosen for its consistent performance in benchmarking studies [10,11] (Fig. 1B). The HLA typing result is then used as a donor-specific reference for expression quantification using *kallisto-bustools* [16]. We tested the accuracy of the allele typing as well as the specificity and accuracy of the subsequent quantification separately to ensure each step’s reliability: accurate allele typing is crucial as it defines the reference for quantification, while accurate quantification is essential for reliable downstream analyses of expression patterns.

To validate both steps, we processed four distinct datasets representing various conditions using *scIGD* (Table 1). These included the AIDA cohort to assess performance across different ethnic backgrounds, the 20k PBMC dataset for whole-transcriptome data, the Multiple Myeloma dataset for amplicon-based data, and the Merkel-cell carcinoma dataset to test compatibility with longitudinal samples. This allowed us to assess *scIGD* across different sequencing platforms and experimental setups.

#### 3.1.1 Validation of the HLA allele typing

To test the HLA allele-typing performance of *scIGD*, we first applied our method to a subset of the AIDA cohort [19] that included 14 scRNA-seq datasets (8 from Korean (KR) and 6 from Japanese (JP) individuals). The allele-typing results revealed common *HLA-A* and *HLA-C* alleles shared across both populations (Fig. 3A; Supplementary Fig. 1A), consistent with their geographical and shared ancestry. For instance, 3 out of the 8 Koreans and 4 out of the 6 Japanese individuals carried the *A*24:02:01:01* allele, which was also overall most abundant in the allele typing results. To further explore the population-wide distribution of this allele, we used the *immunotation* package [26] to determine its frequency across various human reference populations from the Allele Frequency Net Database [27] (Fig. 3B). As anticipated, the *A*24:02* allele group was found to be more frequent in East Asian populations or those of East Asian descent compared to other populations. Similarly, we conducted this analysis for the *C*01:02* allele group, and observed comparable results, with a higher frequency in East Asian populations (Supplementary Fig. 1B).

**Fig. 3.**
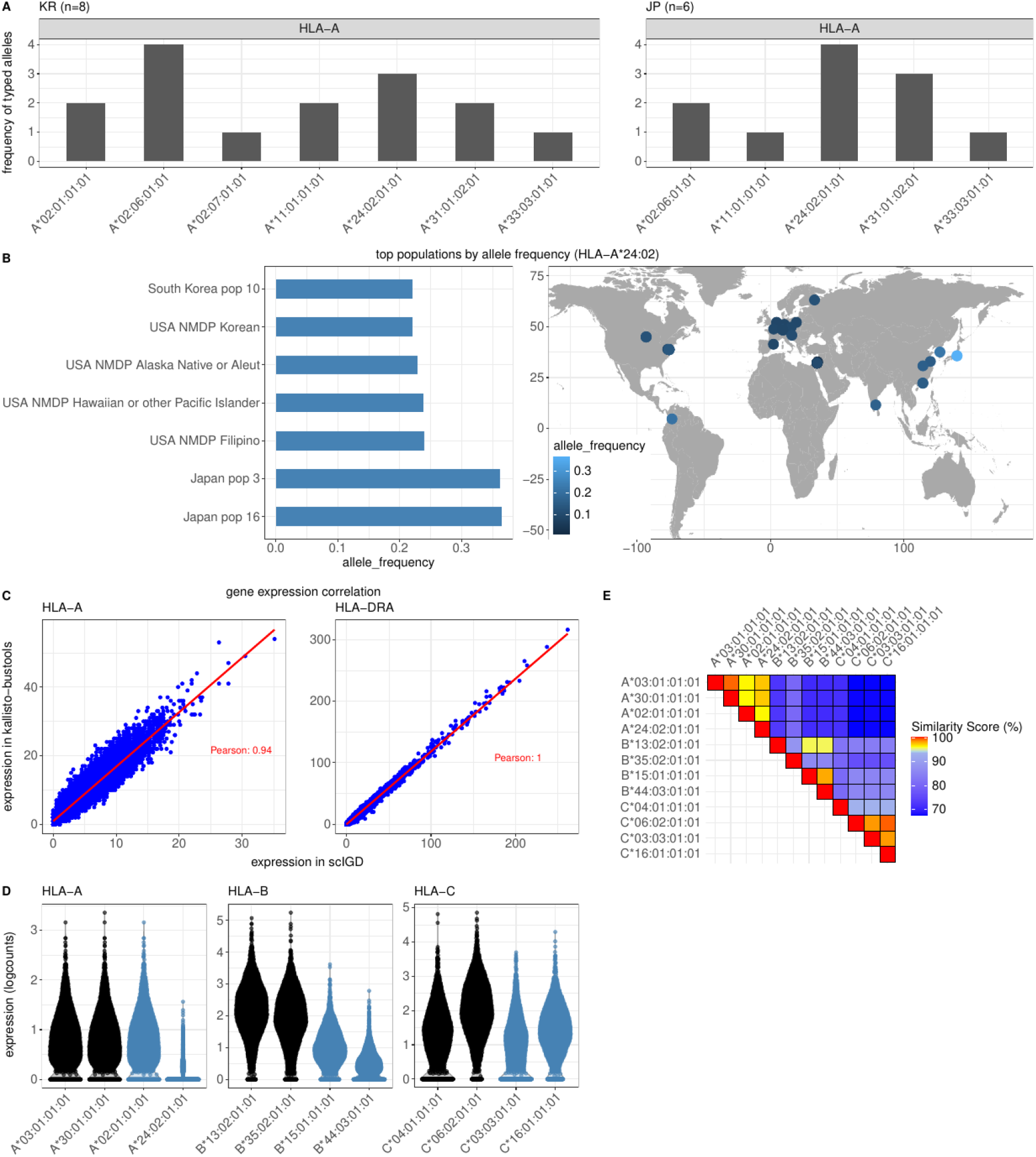
Validation of HLA allele-typing, specificity, and quantification in WTA-based data. **A)** Absolute frequency of *HLA-A* alleles typed in a subset of the AIDA dataset [19], including 14 samples: 8 Korean (KR - left panel) and 6 Japanese (JP - right panel) individuals. HLA allele-typing was performed using the *scIGD* workflow, with no prior information about ethnicity or population. **B)** Relative frequency of the most abundant *HLA-A*24:02* allele group identified in (A) in different human reference populations from the Allele Frequency Net Database (AFND) [27]. The bar graph on the left shows the 7 highest relative frequencies of the *HLA-A*24:02* allele group in AFND. The map on the right shows a visualization of the allele group frequencies in human populations worldwide. **C)** Comparison of *HLA-A* and *HLA-DRA* gene expression quantification using *scIGD* (x-axis) versus kallisto-bustools without using its -mm parameter (y-axis). Raw gene expression counts in the 20k PBMC dataset are shown. HLA gene expression in scIGD is calculated by aggregating the counts of alleles corresponding to each gene. Pearson correlation coefficient is indicated. **D)** Assessment of HLA allele specificity in the Merkel cell carcinoma PBMC pre-treatment dataset. The plot displays normalized expressions of different HLA alleles, with alleles typed by *arcasHLA* [9] in this particular dataset highlighted in black, and those from an unrelated dataset in blue. **E)** A heatmap illustrating the sequence similarity percentages of HLA allele sequences from panel D), calculated with *EMBOSS Needle* [29].

#### 3.1.2 Validation of the HLA expression quantification

Following allele-typing, the *scIGD* workflow quantifies expression levels of all genes and HLA alleles using a donor-specific reference built from the allele typing result. The HLA expression at the gene level is then calculated by aggregating the counts of all alleles belonging to the same HLA gene.

We tested whether this allele aggregation approach for gene-level quantification also correlated with the HLA gene expression levels obtained by conventional quantification methods for 10x Genomics (Fig. 3C) and BD Rhapsody single cell sequencing data (Supplementary Fig. 1C). To assess how our HLA gene counts compare to those obtained using *kallisto-bustools*, where no allele-specific information is provided and gene quantification is based solely on a default reference genome, we compared the raw gene expression counts of *HLA-A* and *HLA-DRA* genes (Fig. 3C) as well as *HLA-C* genes (Supplementary Fig. 1C). The Pearson correlation coefficient was close to 1, indicating that the HLA gene-level expression results derived from *scIGD*’s aggregated allele counts were highly comparable to those from conventional quantification methods. This supports the reproducibility and accuracy of HLA quantification in *scIGD*.

#### 3.1.3 Validation of the specificity of the HLA allele typing

In the quantification step of *scIGD*, we include reads that pseudo-align to multiple transcripts using the *-mm* option of *kallisto-bustools*. This is necessary because the HLA alleles in the donor-specific reference files are very similar in sequence and their expression would otherwise be underestimated.

In order to test the specificity of our quantification approach, we next examined how *scIGD* quantified unrelated HLA alleles that were not identified in HLA allele typing. For this, we quantified expression levels for alleles identified in the HLA typing step in the 20k PBMC dataset (colored in black) alongside unrelated alleles (colored in blue) (Fig. 3D). As hypothesized, the alleles corresponding to the 20k PBMC dataset generally showed higher expression levels compared to external alleles, with exceptions observed for unrelated alleles *HLA-A*02:01* and *HLA-C*16:01*, which exhibited unexpectedly high expression levels.

We further assessed whether these unexpected expression levels could be attributed to sequence similarity. We aligned all allele sequences used in Fig. 3D using *Clustal Omega* for multiple sequence alignment (MSA) [28] (Supplementary File 1). Furthermore, we calculated pairwise sequence similarity percentages with *EMBOSS Needle* [29] (Fig. 3E). Our analysis suggested that high similarity near the poly-A tail of a transcript has the highest influence on quantification specificity (Supplementary File 1).

A similar validation approach was applied to the amplicon-based Multiple Myeloma dataset [22], containing 8 donors. Here, we examined the expression of two distinct *HLA-C* alleles and the *HLA-C* gene across all donors (Supplementary Fig. 1D). *Allele I*, identified in donors D3 and D4, showed notably higher expression levels in these two donors than in the others. Likewise, *Allele II* was primarily expressed in donors D1, D5, and D7. Despite these allele-specific differences, *HLA-C* gene-level expression remained uniform across all donors.

### 3.2 Biological application and downstream analysis using SingleCellAlleleExperiment

After performing HLA allele-typing and expression quantification with *scIGD*, we used the resulting quantifications to showcase how our method can be used to explore different immunological questions. For this, we applied the functionality of *SCAE* and performed downstream analyses on the Merkel-cell carcinoma and 20k PBMC datasets.

*SCAE* offers a multi-layer data framework that allows immune genes to be represented across multiple annotation levels—alleles, genes, and functional classes (Fig. 1C). At the allele level, counts are specific to each individual allele, offering detailed insights into allele-specific expression across cells within one donor. Moving up to the gene level, allele counts are aggregated to reflect the total expression of each HLA gene, giving a broader view of gene expression without focusing on individual alleles and allowing comparisons between donors. Finally, at the functional class level, gene-level counts are further aggregated by functional class, representing overall expression for functional gene groups (e.g. HLA Class I and Class II) that correspond to specific immune functions. This multi-layer design allows users to focus on specific immune gene annotation levels within the same data object during their downstream analyses. *SCAE* integrates smoothly with workflows based on *SingleCellExperiment* (*SCE*) [14], making it user-friendly and compatible with a vast array of existing Bioconductor packages for single-cell analysis, while enabling the exploration of immunogenetic diversity.

#### 3.2.1 Detection of HLA loss in single cell sequencing data from solid tumor

We first demonstrate how the multiple layers can be combined to explore loss of HLA expression in tumors. For this, we used the Merkel-cell carcinoma dataset (PBMC and tumor samples, pre- and post-treatment) for which HLA loss has previously been reported [20]. The original study reported that in tumor samples, post-treatment expression of *HLA-B* was significantly downregulated, suggesting a specific immunologic pressure exerted by the immune system on *HLA-B*, particularly from the *HLA-B*35:02*-restricted CD8+ T cells that were transferred as part of the treatment [20].

After preprocessing, we retained 1794, 3439, 2118, and 4896 cells from PBMC pre- and post-treatment and tumor pre- and post-treatment samples, respectively. All four samples were merged into a single *SCAE* object for integrated analysis. HLA class I expression was highest in the immune cell population of PBMC samples and significantly lower in tumor samples composed mainly by tumor cells (Fig. 4A).

**Fig. 4.**
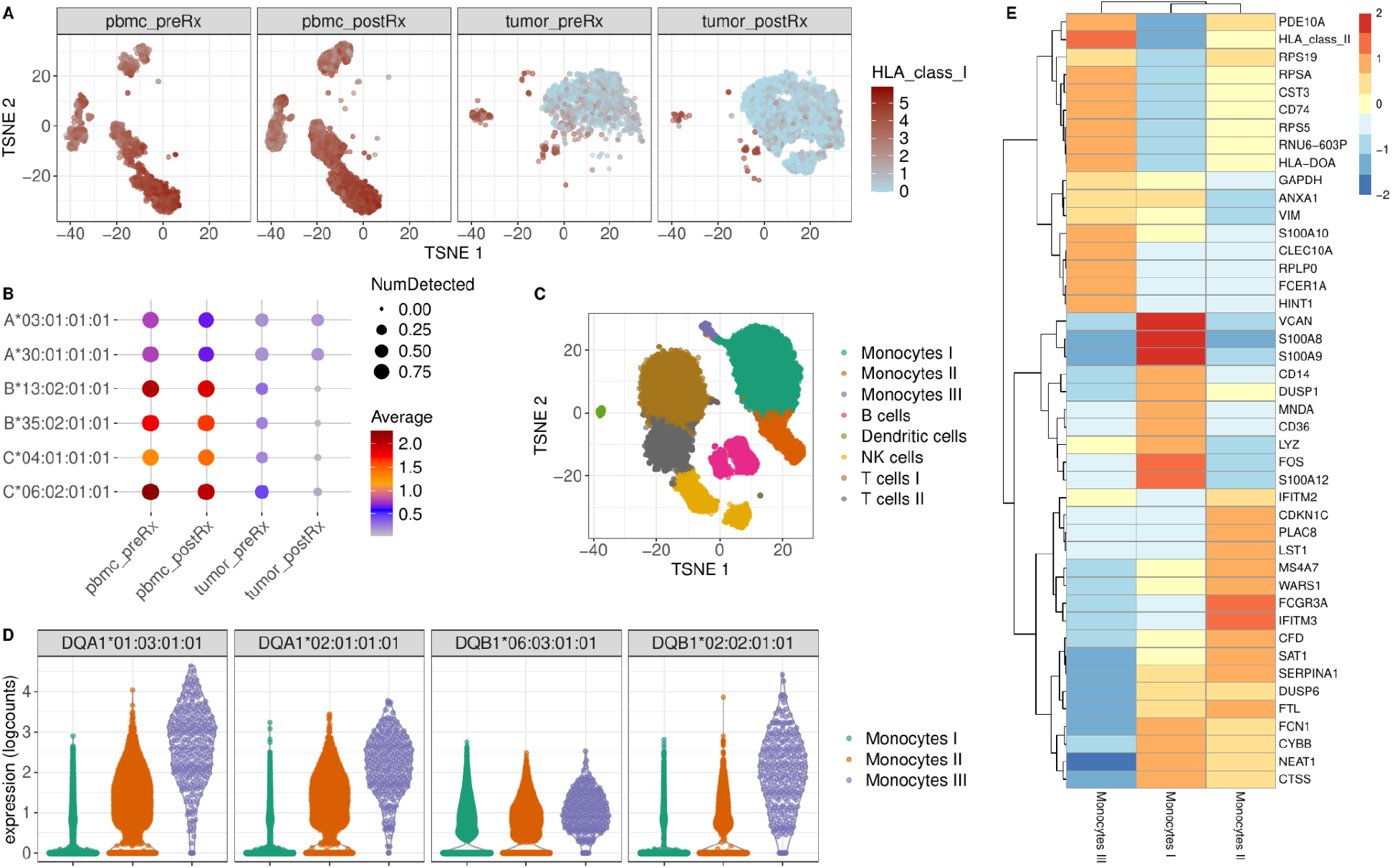
Application of scIGD for biological discovery. **A+B)** Results from the Merkel-cell carcinoma dataset, which includes PBMC (“pbmc_*”) and tumour (“tumor_*”) samples, pre- and post-treatment (“*_preRx” and “*_postRx”), to explore HLA loss in a solid tumor. **A)** Visualization of the log-transformed normalized expression of HLA class I functional gene group using a t-SNE plot. t-SNE was generated from a batch-corrected dimensionality reduction using the MNN integration method and is faceted by sample. Notably, HLA class I expression is high in PBMC samples and significantly lower in tumor samples. **B)** Bubble plot representation of HLA class I allele expression across the four samples from the same Merkel cell carcinoma dataset as in A). The color scale reflects the average of the log-transformed normalized expression values, highlighting differences in allele expression among the samples, while the size of the dots encodes the proportion of cells expressing HLA class I. **C+D+E)** Results from the 20k PBMC dataset to assess HLA allele-specific expression in different cell types and subsets. **C)** Cluster annotation on t-SNE. Louvain clustering was performed using the shared nearest neighbor (SNN) graph constructed from the PCA results. Cluster annotation was achieved using candidate marker genes identified through differential expression testing between pairs of clusters. **D)** Log-transformed normalized expression values of *HLA-DQA1* and *HLA-DQB1* alleles in the three monocyte subclusters from the 20k PBMC dataset. **E)** Heatmap showing the centered mean expression values of the most important marker genes for the three monocyte subclusters.

Our findings align with the original study’s observations, as we also detected a marked reduction in HLA class I gene expression among tumor post-treatment cells, specifically showing a complete loss of both *HLA-B* alleles (Fig. 4B). In addition, our allele typing results correctly identified *HLA-B*35:02*, reinforcing the connection to the selective pressure discussed in the original study. Since we observed downregulation of both *HLA-B* alleles, the mechanism of HLA downregulation is likely not allele-specific in this particular example. In summary, the *SCAE* functionality allowed us to explore loss of HLA expression at both the gene and allele levels, supporting the conclusions drawn in the original study.

#### 3.2.2 Identification of differential allele expression in a subset of antigen presenting cells

Next, we demonstrate the use of our method for detection of differential HLA allele expression in different cell types and subsets. For this, we used the 20k PBMC dataset that can also be found in our *scaeData* R/ExperimentHub package (https://bioconductor.org/packages/scaeData).

After preprocessing, 19765 cells were retained. Cell-type annotation, projected on the *t-SNE*, provided a clear visualization of distinct immune subsets, where clusters representing different cell types were readily distinguishable (Fig. 4C). This included three monocyte subsets, B cells, dendritic cells, NK cells, and two T cell subsets, which were characterized by distinct marker expression profiles (Supplementary Fig. 2).

We further explored the expression of *HLA-DQA1* and *HLA-DQB1* alleles in the three monocyte subsets (Fig. 4D). Monocytes I exhibited the lowest expression levels of *HLA-DQA1* and *HLA-DQB1*, whereas Monocytes III had the highest. In addition, we observed differential *HLA-DQB1* allele expression profiles between the subsets: Monocytes I and II demonstrated a higher expression of the *HLA-DQB1*06:03*, while Monocytes III favored *HLA-DQB1*02:02*, suggesting differences in antigen-presenting function.

The overall gene expression profiles of the different monocyte populations indeed also revealed functional heterogeneity (Fig. 4E). Monocytes I upregulated classical pro-inflammatory markers, such as *S100A8, S100A9, VCAN, FOS, LYZ*, and *CD14*, indicative of a role in acute inflammation. In contrast, Monocytes III showed increased expression of genes associated with HLA class II molecules and other markers like *PDE10A* and *HINT1*, suggesting they might be non-classical, serving a more specialized role in antigen presentation or immune regulation.

These findings emphasize how allele-specific HLA expression can delineate functionally distinct subsets within the same cell type, revealing their distinct biological roles. This level of allele-specific resolution, detecting differential HLA allele expression across subsets of the same cell type, is a significant advantage of our method over traditional transcriptomic workflows, which focus on gene-level expression alone. This highlights the capacity of our workflow to detect variation in HLA allele expression even within subsets of the same cell type.

## 4 DISCUSSION

In this study, we present a comprehensive workflow that enables allele-specific analysis of HLA expression in single-cell RNA sequencing (scRNA-seq) data. By leveraging *scIGD* and *SingleCellAlleleExperiment* (*SCAE*), we demonstrate the utility of our tools across diverse scRNA-seq datasets, including whole-transcriptome 10x Genomics datasets and a multiplexed amplicon-based BD Rhapsody dataset. Our results highlight the ability of this workflow to not only accurately perform HLA allele-typing but also quantify allele-specific expression, offering novel insights into human immunogenomic diversity.

We validated the allele-typing results across multiple datasets and demonstrated the robustness and specificity of the workflow (Fig. 1A, B, D; Supplementary Fig. 1A, B, D). The strong correlation observed between gene-level quantification from our method and standard tools, such as *kallisto-bustools* [16], further supports the accuracy of our allele-specific quantification approach (Fig. 1C; Supplementary Fig. 1C). Across both whole-transcriptome and amplicon-based datasets, our method was able to reliably detect allele-specific expression differences. Moreover, our methodology reliably recapitulates findings from previous studies such as the loss of *HLA-B* expression in a Merkel-cell carcinoma case (Fig. 4B).

As an advancement over existing tools such as *seq2HLA [5], OptiType* [6], *PHLAT* [7], *arcasHLA* [9], and *scHLAcount* [12], our method integrates both typing and allele-specific quantification within a single streamlined workflow, and at the same time offers a data structure optimized for interactive single-cell downstream analysis. Of note, *SCAE* can be integrated with interactive data exploration tools such as *iSEE* [30]. One of the key assets of our analysis workflow is the capacity to detect HLA expression differences across subsets of the same immune cell type, particularly within the monocyte subsets in the 20k PBMC dataset (Fig. 4D, E). This type of allele-specific expression difference is not detectable with traditional transcriptomic workflows [14,31,32]. By capturing such subtle variations, our method opens new avenues for exploring immune cell heterogeneity and the functional implications of HLA expression.

Our study highlights the importance of integrating allele-level information into scRNA-seq analyses, particularly for important and structurally diverse immune mediators such as the HLA molecules. Future research could build on this work by extending our workflow to other highly polymorphic loci such as B and T cell receptors, and killer Ig-like receptors (KIRs). Overall, the *scIGD* workflow for allele-specific quantification in scRNA-seq offers a powerful tool for exploring the expression of important immune mediators across human donors at very high molecular detail.

## Data availability

AIDA: The entire AIDA cohort FASTQ files are available via the Human Cell Atlas Data Coordination Platform [https://data.humancellatlas.org/explore/projects/f0f89c14-7460-4bab-9d42-22228a91f185]. The donor IDs of the 14 datasets analyzed were the following: KR_SGI_H022, KR_SGI_H025, KR_SGI_H040, KR_SGI_H045, KR_SGI_H049, KR_SGI_H095, KR_SGI_H098, KR_SGI_H102, JP_RIK_H016, JP_RIK_H035, JP_RIK_H047, JP_RIK_H056, JP_RIK_H074, and JP_RIK_H148. Reference: Kock KH, Le Min T, Han KY *et al*. Single-cell analysis of human diversity in circulating immune cells. *bioRxiv* 2024:2024.06.30.601119.

Merkel-cell carcinoma: The FASTQ files are available via the National Center for Biotechnology Information Gene Expression Omnibus (NCBI GEO), accession GSE 117988 [https://www.ncbi.nlm.nih.gov/geo/query/acc.cgi?acc=GSE117988]. The samples analyzed were the following: GSM3330561, GSM3330564, GSM3330559, and GSM3330560. Reference: Paulson KG, Voillet V, McAfee MS *et al*. Acquired cancer resistance to combination immunotherapy from transcriptional loss of class I HLA. *Nature Communications* 2018;9:1–10.

20k PBMC: The FASTQ files are available via the 10x Genomics website [https://www.10xgenomics.com/datasets]. Dataset name: 20k Human PBMCs, 3’ HT v3.1, Chromium X. Reference: Datasets. *10x Genomics*.

Multiple Myeloma: Reference: Enssle JC, Campe J, Moter A *et al*. Cytokine-responsive T- and NK-cells portray SARS-CoV-2 vaccine-responders and infection in multiple myeloma patients.

*Leukemia* 2023;38:168–80. The FASTQ files were made available via the lead author of the original study.

The output of scIGD for all datasets, including the count matrices, is available via Zenodo [https://doi.org/10.5281/zenodo.14033960].

## Code availability

All code used to analyze the four datasets and generate the figures is available in the following GitHub repository: https://github.com/AGImkeller/scIGD_manuscript_2024.

*scIGD* is available only on GitHub (https://github.com/AGImkeller/scIGD), whereas *SingleCellAlleleExperiment* and *scaeData* are available on GitHub and Bioconductor (https://bioconductor.org/packages/SingleCellAlleleExperiment and https://bioconductor.org/packages/scaeData). All three tools are available under the MIT license.

## Acknowledgements

The authors thank Aakanksha Singh and Samira Ortega Iannazzo for valuable feedback on the manuscript. The authors would like to thank Evelyn Ullrich, Ivana von Metzler, Julius Enssle, and Michael Rieger for providing the FASTQ files of the multiple myeloma dataset and for technical advice on the sequencing technologies. The authors would like to thank the Asian Immune Diversity Atlas (AIDA) team for making the respective FASTQ files available.

## Supplementary data

**Supplementary Fig. 1.**
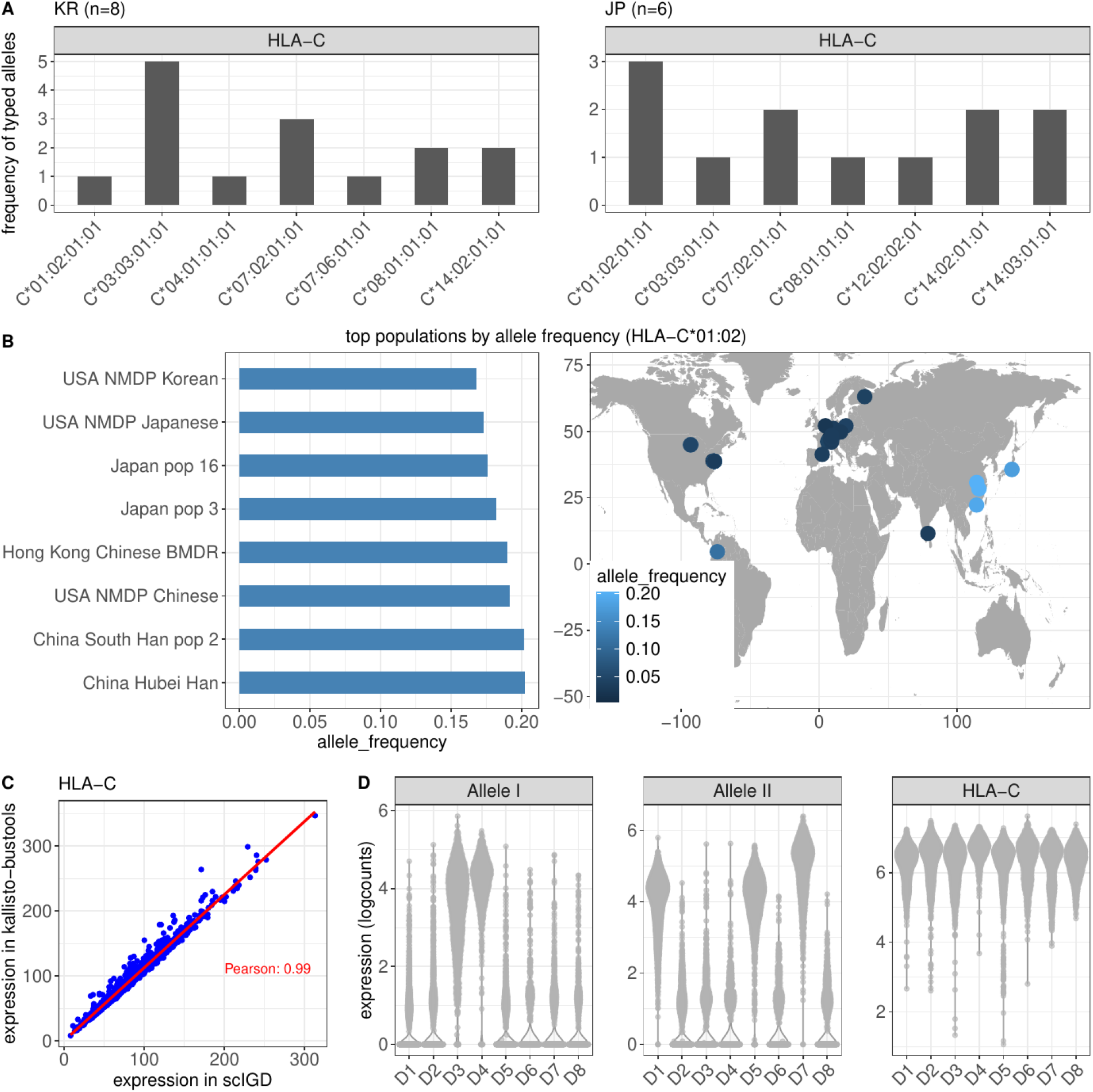
Validation of HLA allele-typing, specificity, and quantification in amplicon-based data. **A)** Absolute frequency of *HLA-C* alleles typed in a subset of the AIDA dataset [19], including 14 samples: 8 Korean (KR - left panel) and 6 Japanese (JP - right panel) individuals. HLA allele-typing was performed using the scIGD workflow, with no prior information about ethnicity or population. **B)** Relative frequency of the *HLA-C*01:02* allele group identified in (A) in different human reference populations from the Allele Frequency Net Database (AFND) [27]. The bar graph on the left shows the 8 highest relative frequencies in AFND. The map on the right shows a visualization of the allele group frequencies in human populations worldwide. **C)** Comparison of *HLA-C* gene expression quantification using scIGD (x-axis) versus kallisto-bustools (y-axis). Raw gene expression counts in the D1 sample of the Multiple Myeloma dataset are shown. HLA gene expression in scIGD is calculated by aggregating the counts of alleles corresponding to each gene. Pearson correlation coefficient is indicated. **D)** Assessment of HLA allele specificity in the Multiple Myeloma dataset, containing 8 donors. The plot displays normalized expressions of two different *HLA-C* alleles and the *HLA-C* gene across all donors. Alleles identified in specific donors exhibited notably higher expression levels compared to those in other donors. In contrast, the expression of the *HLA-C* gene remained uniform across all donors. Note: Only two alleles are shown here for illustrative purposes, but additional alleles were also typed from these 8 donors.

**Supplementary Fig. 2.**
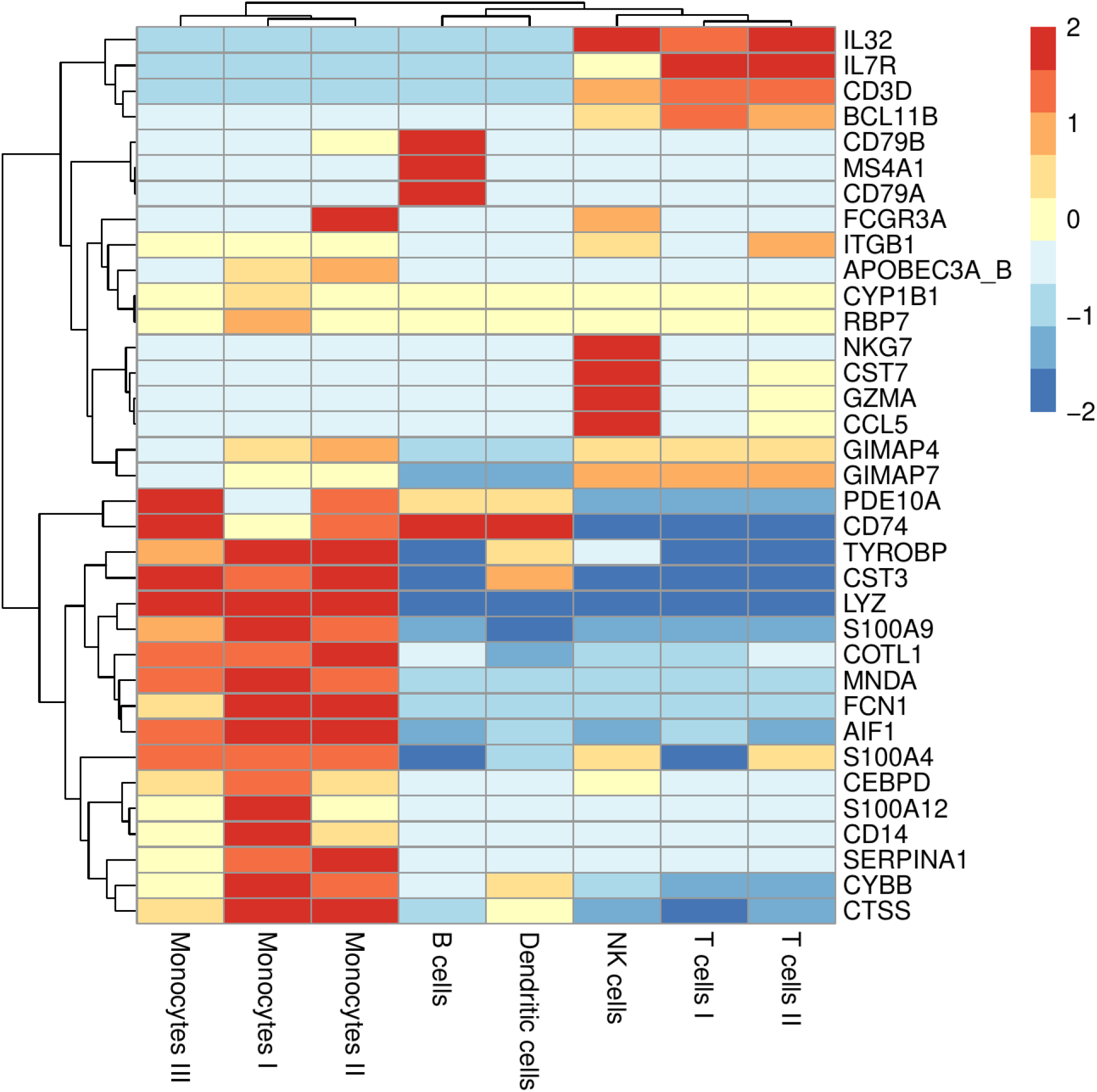
Gene expression profiling for cluster annotation in 20k PBMC dataset. Heatmap showing the centered mean expression values of the most important marker genes for the different single-cell clusters in the 20k PBMC dataset. Cluster annotation was achieved using these marker genes.

