## Supplementary figures and images for "A comprehensive workflow for allele-specific immune gene quantification and expression analysis in single-cell RNA-seq data"

### SupplementaryFigure1.pdf

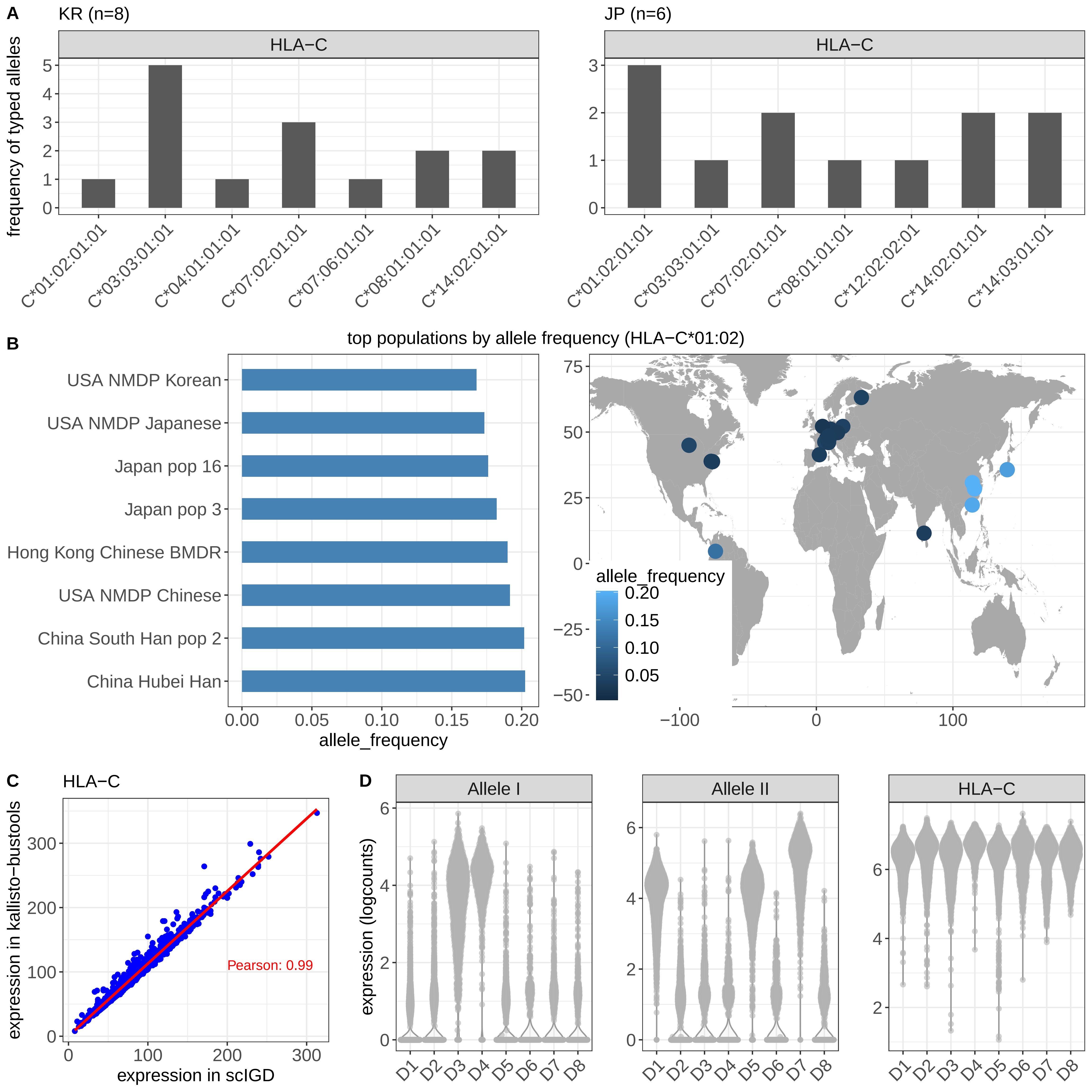

### SupplementaryFigure2.pdf

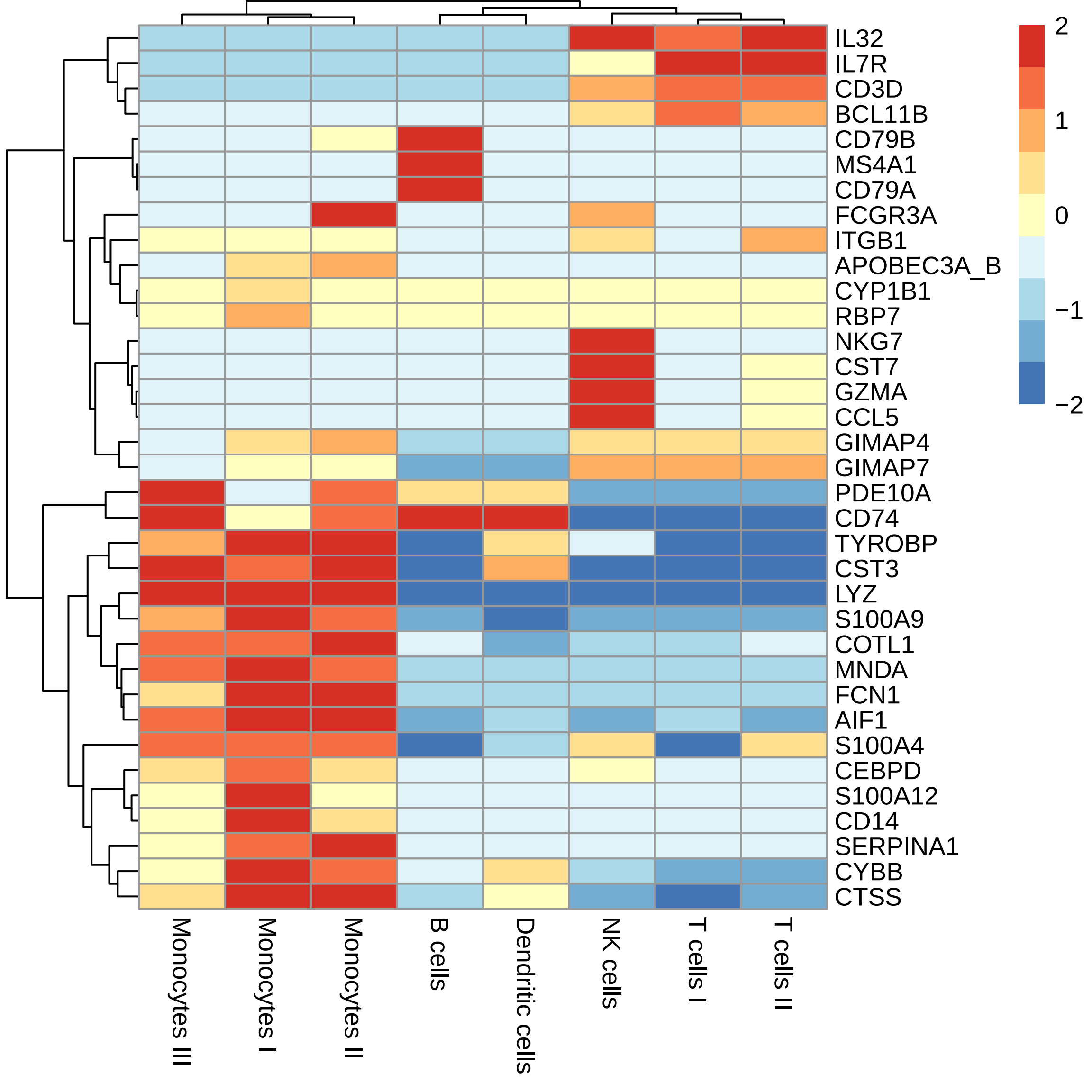
